# Treatment of canine visceral leishmaniasis with Milteforan^™^ induces *Leishmania infantum* resistance to miltefosine and amphotericin B

**DOI:** 10.1101/2021.04.08.438938

**Authors:** Gustavo Gonçalves, Monique Paiva Campos, Alessandra Silva Gonçalves, Lia Carolina Soares Medeiros, Fabiano Borges Figueiredo

## Abstract

Visceral leishmaniasis (VL) is the most severe form of leishmaniasis and is caused by *Leishmania infantum* in the Americas. Since the use of Milteforam™ was authorized to treat canine visceral leishmaniasis (CVL) in Brazil in 2017, there has also been fear of the emergence of parasites resistant to this drug and, through cross-resistance mechanisms, to meglumine antimoniate and amphotericin B. Additionally, the literature shows that acquisition of resistance is followed by increased parasite fitness, with higher rates of proliferation, infectivity and metacyclogenesis, which are determining factors for parasite virulence. In this context, this study aims to analyze the impact of treating a dog with Milteforan™ on the generation of parasites resistant to miltefosine, meglumine antimoniate, and amphotericin B. To this end, *in vitro* susceptibility tests were conducted against these drugs with T0 (parasites isolated from the dog before treatment with Milteforan™), T1 (after one course of treatment), and T2 (after two courses of treatment) isolates. The rates of cell proliferation, infectivity, and metacyclogenesis of the isolates were also evaluated. The results indicate a gradual increase in parasite resistance to miltefosine and amphotericin B with increasing the number of treatment courses. A trend increase in the metacyclogenesis rate of the parasites was also observed as drug resistance increased. Therefore, treatment of CVL with Milteforan™ induces resistance to miltefosine and amphotericin B as well as changes in parasite fitness, and may have an impact on animal and human public health.

## Introduction

Leishmaniasis is a parasitic disease caused by protozoa of the genus *Leishmania* and transmitted by the bite of infected sandflies. Its most severe form, visceral leishmaniasis (VL) (1), shows clinical characteristics of severe evolution in humans (2), with occurrence of a zoonotic cycle both in South America and in the Mediterranean Sea region (3,4). In this context, *Leishmania infantum* (syn = *Leishmania chagasi*) is the most important etiological agent involved (5) and the dog (*Canis familiaris*) is the main host (2). Canine leishmaniasis preceded the occurrence of human cases (2,6,7), and control of the zoonotic cycle is still a challenge (3).

Treatment options for leishmaniasis are limited and unsatisfactory. For more than 60 years, human treatment was centered on the use of pentavalent antimonials (8). After that, new drugs were developed, but the arsenal to treat VL is still limited, thus characterizing this disease as neglected. Current therapies rely on three main drugs: pentavalent antimonials (first drug of choice), amphotericin B, and miltefosine (9).

Treatment with miltefosine presents important limitations because of its teratogenic character and long half-life, which facilitate the emergence of parasite resistance that can be easily established through isolated point mutations (10). Additionally, monotherapy based on this drug is no longer recommended in humans because of the observed flaws and the rapid acquisition of drug resistance by the parasites (11,12), and it should now be used in combination with other anti-*Leishmania* drugs (9).

In 2017, the use of Milteforan™ was authorized to treat canine visceral leishmaniasis (CVL) in Brazil (13). However, treatment failures have been observed both in monotherapy and combined therapies (14,15), with improvement in canine symptomatology not followed by parasitological clearance (16). Thus, treatment is still not considered an effective control measure because, in addition to the risk of parasite resistance, relapses are frequent and dogs can continue to infect the invertebrate host even weeks after the end of treatment, despite being clinically cured (17).

Therefore, in addition to the risk involving the treatment of dogs with miltefosine in relation to the possible emergence of parasites resistant to this drug, there are reports of cross-resistance to other drugs (18–23), which can lead to the emergence of parasites resistant not only to miltefosine, but also to other drugs used to treat CVL, further aggravating the public health problem, especially after the authorization of this treatment in dogs in 2017 (13).

Moreover, there are also reports on the impact of acquisition of resistance on parasite fitness, where drug-resistant parasites presented higher rates of cell proliferation, metacyclogenesis, and infectivity compared with those of susceptible parasites(24–26), which are aggravating factors of disease virulence (27) that may have an impact on public health.

In this context, the present study aimed to analyze the impact of treating a dog with CVL with Milteforan™ on the generation of parasites resistant to miltefosine, as well as to meglumine antimoniate and amphotericin B, through cross-resistance mechanisms. It also aimed to determine the impact of possible acquisition of resistance on the rates of cell proliferation, metacyclogenesis, and infectivity of the parasite.

## Methods

## Experimental design and collection of isolates

The isolates used in this study were collected from a naturally infected, mixed-breed female dog, aged approximately 5 years, from the municipality of Campo Grande, state of Mato Grosso do Sul, Brazil. After positive serological diagnosis using the Dual-path Platform chromatographic immunoassay (DPP^®^), additional collections were performed to confirm the infection by *Leishmania infantum* through quantitative Polymerase Chain Reaction (qPCR) and parasitological culture. For the qPCR, a 3 mm diameter intact skin fragment was obtained by punch biopsy and stored in a sterile flask free of RNase and DNase at –20 °C. For the parasitological culture, in addition to another skin fragment, bone marrow and lymph node aspirates were collected and stored in sterile saline solution containing antibiotics and antifungals under refrigeration. The samples were kept at 4 °C for 24 h, sown in biphasic culture medium containing Novy-MacNeal-Nicole (NNN) medium and Schneider’s insect medium supplemented with 10% fetal bovine serum (FBS), and examined weekly by optical microscopy in search of promastigote forms of the parasite for one month (28). Confirmation of infection and characterization of the parasite as *L. infantum* was performed using qPCR with specific species primers. After DNA extraction, the sample was amplified using the TaqMan^®^ system on the StepOne™ platform (Applied Biosystems^®^). The TaqMan^®^ MGB probe and the qPCR were designed to target the conserved regions of the *L. infantum* KDNA. The primers LEISH-1 (5’-AACTTTTCTGGTCCTCCGGGTAG-3 ‘) and LEISH-2 (5’-ACCCCCAGTT TCCCGCC-3’) and the probe TaqMan-MGB (FAM-5’AAAAATGGGTGCAGAAAT-3’-NFQM-3GB) (29,30) were used. The samples were amplified on the StepOne™ platform. After confirmation of infection with by all proposed methodologies (DPP^®^, qPCR, and parasitological culture), treatment with Milteforan™ was started. The treatment was carried out according to the manufacturer’s instructions in two courses with an interval of four months between them. In each treatment course, 20 mg/kg of the drug was administered in daily doses for 28 consecutive days. New collections were performed immediately before the start of the new course aiming to isolate, in addition to the parasites already isolated prior to treatment commencement (T0), parasites after one (T1) and two (T2) courses of treatment, as shown in Figure 1. The study was approved by the Ethics Committee on Animal Use of FIOCRUZ under protocol no. P-12 / 2020-6.

**Figure 1:**
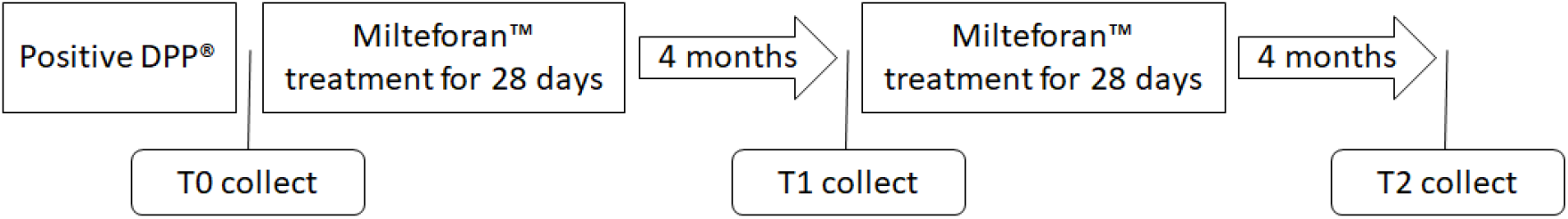
Schematic representation of the experimental design consisting of collection of T0 (parasites collected immediately after positivity in the DPP^®^ test and before treating the dog with Milteforan™) isolates followed by treatment of the dog with the drug for 28 consecutive days; after four months, collection of T1 (parasites collected from the dog after one course of treatment) isolates followed by another treatment of the dog with Milteforan™ for 28 days; finally, after another 4-month interval, collection of T2 (parasites collected from the dog after two courses of treatment) isolates.

### Drugs

The commercial drugs Milteforan™ (Virbac^®^), Glucantime (generic pharmacy), and Amphotericin B (generic pharmacy) were used for the *in vitro* assays as a sources of miltefosine, meglumine antimoniate, and amphotericin B, respectively. The drugs were stored as indicated on their package inserts and diluted immediately before the assays in Schneider’s culture medium until the desired concentrations were reached.

### *In vitro* susceptibility of isolates to miltefosine, meglumine antimoniate, and amphotericin B

Aiming to evaluate the possible emergence of drug resistant parasites as a result of treating the dog with Milteforam™, the half maximal inhibitory concentration (IC_50_) values of the *L. infantum* T0, T1, T2, and reference strain (MHOM/BR/74/PP75) promastigote and amastigote forms were determined against miltefosine, meglumine antimoniate, and amphotericin B. The IC_50_ values against promastigote forms were determined using the MTT (3-(4,5-Dimethylthiazol-2-yl)-2,5-Diphenyltetrazolium Bromide) colorimetric assay. Promastigote cultures in exponential growth were adjusted to a concentration of 1×10^6^ parasites/mL and incubated (25 °C) in 96-well plates (200 µL per well) with the drugs at different concentrations for 24 h. After incubation, the cultures had their viability determined with addition of 20 µL MTT (5 mg/mL) to each well of the plate, followed by incubation at 36 °C for 3 h and solubilization of the formate crystals with 20 µL 10% sodium dodecyl sulfate (SDS) and 30 µL 100% dimethyl sulfoxide (DMSO). Absorbance reading of the wells was performed on a spectrophotometer at 550 nm (31). A control (no drugs added) was used for each isolate. The IC_50_ values were determined using cell viability values for each concentration of each drug.

For assays against the amastigote forms of the isolates and the reference strain, the THP-1 cell line was used as a host. The monocytes were kept at 37 °C in a humid incubator, under an atmosphere of 5% CO_2_, in RPMI 1640 medium supplemented with 10% FBS, HEPES, and 1% antibiotic (Penicillin Streptomycin, Sigma). Cultures were maintained by weekly breeding until their growth reached 1×10^6^ cells/mL. Thereafter, THP-1 cells were seeded in 96-well plates at a density of 51×10^4^ cells/well in RPMI 1640 medium containing 200 nM phorbol myristate acetate (PMA). The plates were then incubated for 96 h to allow cell differentiation to adhered macrophages, and the culture medium was replaced with a new one without PMA after 48 h. Concomitantly with this process, the isolates and the reference strain of *L. infantum* were cultured up to 6-7 days in order to be able to inoculate cells already adhered and differentiated in the macrophages. Inoculation was carried out in the proportion of 10 parasites per cell (10:1) and incubated in wells containing the differentiated cells for 4 h. The different drug concentrations were then added (in triplicate per evaluated dose) to each well and the plates were incubated for 48 h. After treatment, the cells were fixed with methanol and stained with DAPI to perform the intracellular amastigote count. A negative control (without treatment) was used as a 100% infection. Inhibitory activity was assessed by counting the number of intracellular amastigotes in 100 cells randomly captured from each well (40x objective). Values were expressed as percentage of inhibition: PI = 100 - ((Tx100) / C), where *T* corresponds to the average number of amastigotes treated and *C* to the average number of amastigotes from the negative control (32). The IC_50_ values were determined using PI values for each concentration of each drug.

### Impact of treatment with Milteforam™on the growth curves and infectivity and metacyclogenesis rates of the isolates

The impact of the possible acquisition of resistance on the fitness, growth curves, and infectivity and metacyclogenesis rates of the parasites was determined. To measure the growth curve, a culture containing the isolates and the reference strain in exponential growth phase was adjusted to the concentration of 1×10^6^ parasites/mL and seeded in a 24-well plate (1 mL per well). The absorbance values were measured at 800 nm every 24 h, for 8 days to correlate the increase in absorbance with the concentration of parasites in the culture (33). For determination of the infectivity rates, THP-1 cells were infected with the isolates and the reference strain as previously described. After fixing, staining, and counting of 100 cells, the average number of amastigotes per cell infected with the isolates and with the reference strain were compared. The metacyclogenesis rates were determined by the negative selection methodology with peanut agglutinin (PNA) (Sigma, St. Louis, MO, USA) (34,35). Briefly, 6-7 day cultures of the isolates and the reference strain were collected by centrifugation at 2000 xg for 10 min and resuspended at a concentration of 2×10^8^ parasites/mL in 10 mL of Schneider’s medium supplemented with 50 μg/mL PNA. The promastigotes were left at room temperature for 30 min for agglutination. Immediately after that, the supernatant and the pellet were collected. The pellet was resuspended in the same initial Schneider’s medium volume with 50 μg/ml PNA. The two fractions were collected by centrifugation at 200 xg for 10 min and the supernatant resulting from both was centrifuged at 2000 xg for 10 min to obtain the metacyclic promastigotes. The number of metacyclic promastigotes was determined by counting in a Neubauer chamber and the percentage of metacyclogenesis among the isolates was calculated by the ratio of the number of metacyclic promastigotes to the total initial promastigote population. All experiments were carried out in triplicate.

### Statistical analysis

Data normality was assessed by a Kolmogorov-Smirnov test and the IC_50_ values was obtained with The GraphPad 5.0 (Prism) program using non-linear regression. To compare all groups was used the parametric one-way analysis of variance (ANOVA), followed by the Tukey test.

## Results

After a positive result in the DPP^®^ test, skin fragments and bone marrow and lymph node aspirates were collected aiming to confirm infection by *L. infantum* and isolate the parasite in culture. The species specific primers used in the qPCR successfully confirmed infection by *L. infantum*, and the parasite was isolated in culture. All tests were repeated immediately before the start of a new treatment course, resulting in three different isolates: MCAN/BR/19/CG06T0 (T0), MCAN/BR/19/CG06T1 (T1), and MCAN/BR/20/CG06T2 (T2), which enabled access to the parasites at different stages throughout the dog’s treatment with Milteforan™, as shown in Figure 1.

### Susceptibility assay

Susceptibility assays conducted with the isolates and the reference strain showed significant increase in the IC_50_ values of the promastigote (Figure 2; Graphs A and C) and amastigote (Figure 2; Graphs B and D) forms of the parasite, thus evidencing resistance to miltefosine and amphotericin B as the number of treatment courses increased. The parasites isolated prior to treatment (T0) presented IC_50_ values against miltefosine equal to that of the control; however, these values increased after only one course of treatment (T1) (Graphs A and B), diverging statistically from the control with the reference strain. The upward trend continued throughout the treatment, with T2 isolates presenting IC_50_ values approximately two times higher compared with that of the reference strain.

**Figure 2:**
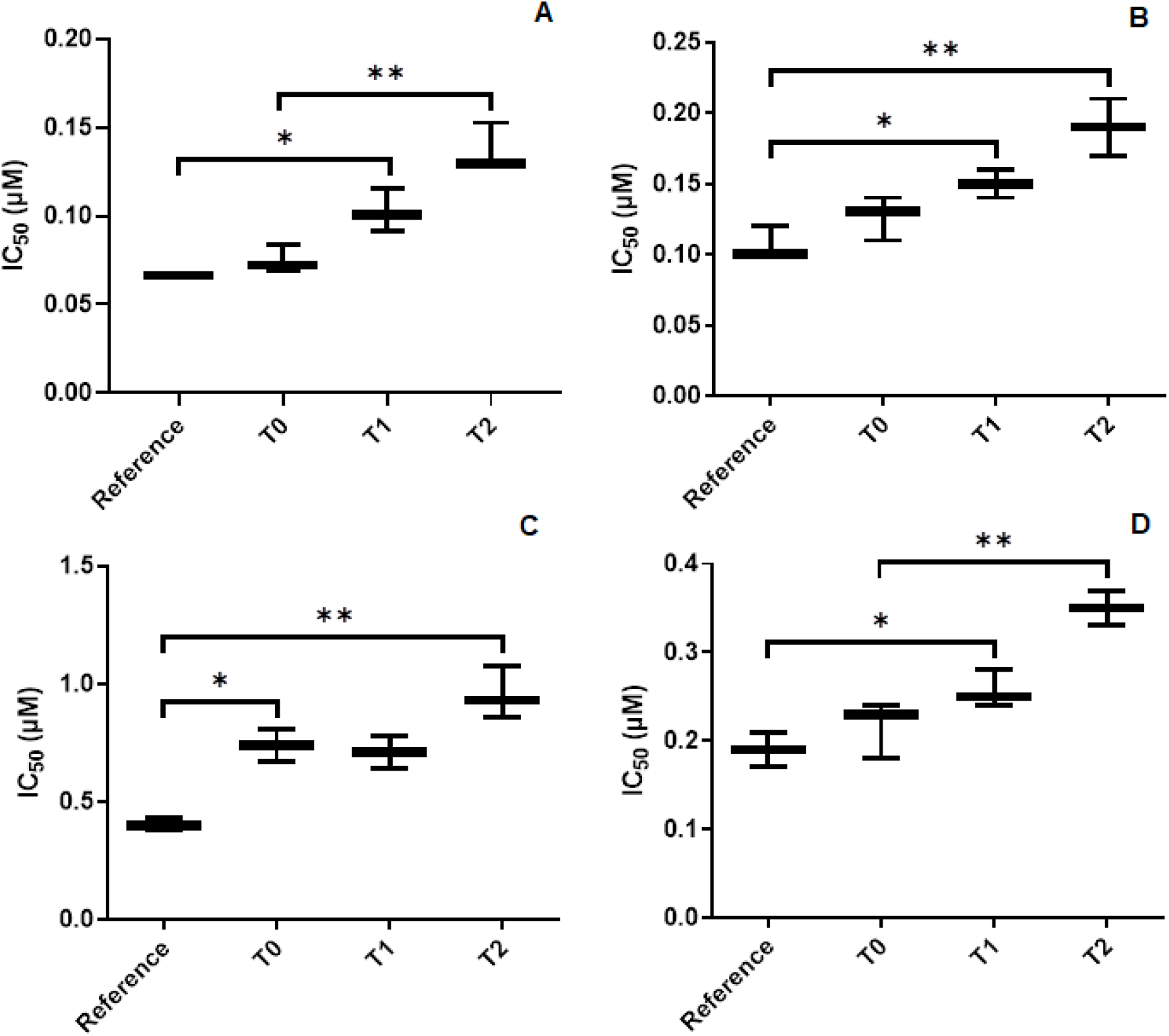
Results of *in vitro* resistance tests of the isolates and the reference strain to the drugs tested. Graphs A and B show the IC_50_ values of the promastigote (Graph A) and amastigote (Graph B) forms of the isolates and reference strain against miltefosine. Graphs C and D show the IC_50_ values of promastigote (Graph C) and amastigote (Graph D) forms of the isolates and reference strain against amphotericin B. Statistical significance is demonstrated by the asterisks, with one asterisk (*) denoting significance <0.05% and two asterisks (**) corresponding to significance <0.01%.

The same pattern was observed in the parasites treated with amphotericin B, where an increase in resistance to the drug was verified throughout the treatment courses (Figure 2; Graphs C and D). In the promastigote forms of the parasites (Graph C), the IC_50_ values of the isolates before the dog’s treatment with Milteforan™ (T0) was already higher than that of the reference strain; this was also a fact for T1 in the assays with amastigote forms (Graph D). No statistical difference was found between the IC_50_ values of the T0, T1, and T2 isolates against meglumine antimoniate; therefore, they did not increase their resistance to this drug as the treatment course with Milteforan™ progressed in the dog.

### Impact of possible acquisition of resistance on the proliferation, metacyclogenesis and infectivity rates of the parasites

Results of the growth curve of the isolates in culture medium showed no difference between the number of parasites or cell proliferation rate on any of the days analyzed. However, when compared with the growth curve of the reference strain, a clear discrepancy in the proliferation rate can be observed after the third day of cultivation (Figure 3). Another difference observed was the time required to reach the growth curve plateau: 5-6 days for the isolates and 6-7 days for the reference strain; in addition, the reference culture showed a parasite density approximately 2.5 five times higher than those of the isolated parasites on its plateau.

**Figure 3:**
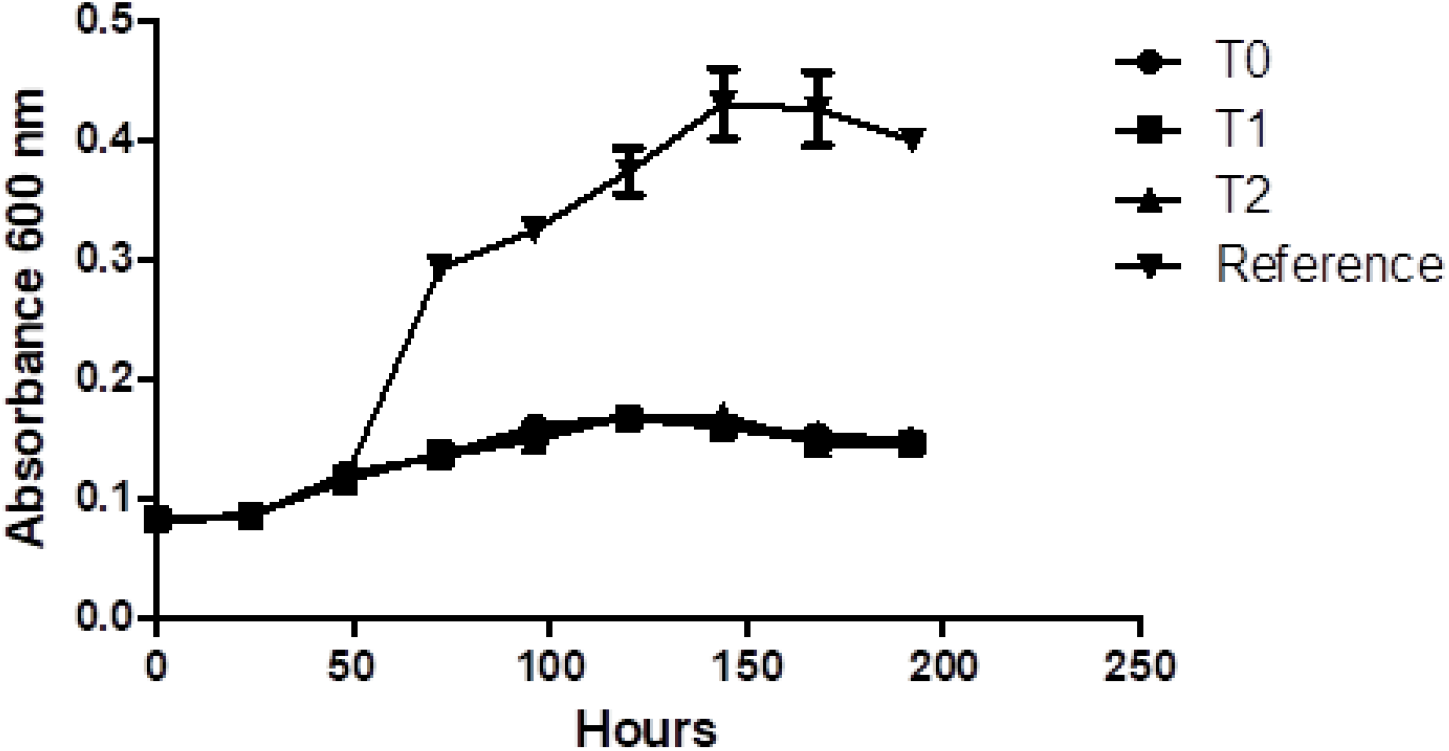
Growth curve of the T0, T1, T2, and reference strain (MHOM/BR/74/PP75) isolates in daily readings of parasite density on a spectrophotometer at 600 nm for eight days.

The impact of acquisition of resistance on parasite infectivity was evaluated by comparing the average number of amastigotes per cell infected with the isolates and with the reference strain. Figure 4 shows that the cells infected with the reference strain have an average of six amastigotes per cell, a number larger than the T0 (fewer than four amastigotes per cell) and T1 and T2 (both with fewer than two amastigotes per cell) isolates.

**Figure 4:**
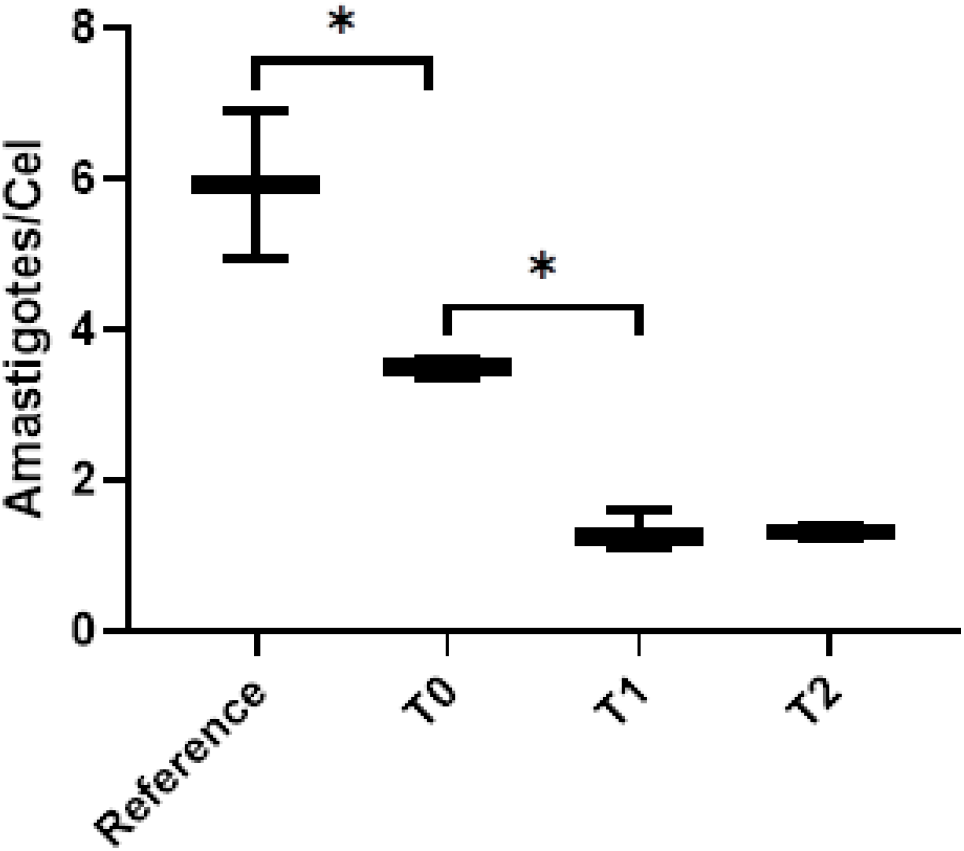
Average number of amastigotes per THP-1 cell infected with the reference strain (MHOM/BR/74/PP75), parasites isolated from the dog before treatment (T0), after one course of treatment (T1), and after two courses of treatment (T2). Statistical significance is shown by asterisks, with one asterisk (*) denoting significance <0.05%.

Analysis of the metacyclogenesis rate of the isolates through differential selection with PNA revealed a tendency of increasing the number of metacyclic promastigotes in the cultures as the number of treatment courses increased. Parasites isolated from the dog before treatment with Milteforan™ showed an average of 2×10^4^ parasites/mL. After one course of treatment, this number increased to about 3×10^4^ parasites/mL, and it reached approximately 5×10^4^ parasites/mL after two treatment courses. Although no statistical difference in the number of metacyclic promastigotes was observed between the isolates at the different stages of treatment, the metacyclogenesis rates show a clear upward trend (Figure 5).

**Figure 5:**
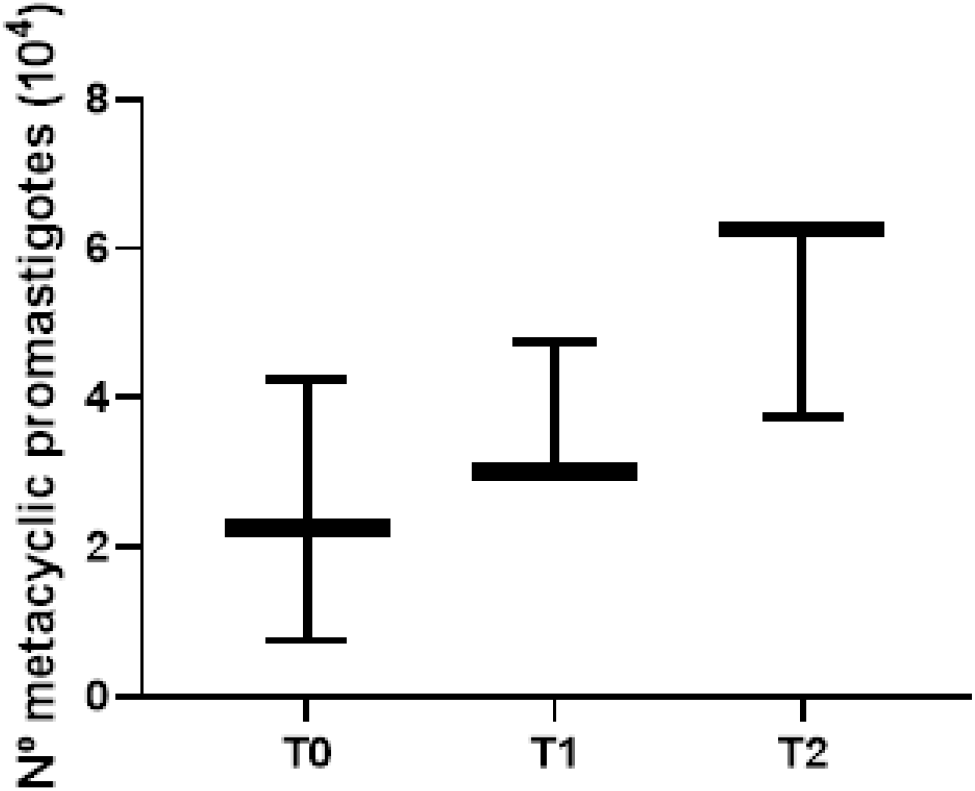
Number of metacyclic promastigotes for each isolate after 6 days of culture and differential selection with peanut agglutinin (PNA).

## Discussion

A strong linear correlation (*R*^2^ = 0.87) was observed between the number of treatment courses with Milteforan™ to which the dog was subjected and the increase in the IC_50_ values of *L. infantum* isolates against this drug. This finding was already expected, considering that the literature brings many reports on the rapid acquisition of resistance by parasites in places where Milteforan™ therapy was adopted (11,12). The results of our experiments can be explained by the fact that the treatment of dogs with Milteforan™ is not considered a totally effective control measure (17), with decreased symptoms not followed by parasitological clearance (16). These parasites that remain in the animal’s organism are exposed to subtherapeutic doses resulting from the long half-life of this drug (10), which produce a selection of resistant parasites and, consequently, induce resistance in the general population in the long term. This *in vivo* dynamic is very similar to that observed in *in vitro* resistance induction experiments (36), where the parasite is exposed to sub-effective and increasing doses of the drug, leading to a gradual increase in its resistance to it. The same behavior has already been observed in the treatment of dogs with meglumine antimoniate, where the larger the number of courses of drug treatment in dogs, the greater the resistance of isolated parasites to this drug (37–39).

It is well established that isolates with lower *in vitro* IC_50_ values are related to cases in which drug therapy is more successful in the clinic (40), with the opposite being also valid. The IC_50_ values of the isolates of the present study at T0 were already higher than the limit proposed by Carnielli et. al. (2019), who established the maximum value for therapeutic success at 0.8 µM (isolates that presented values above this limit were often associated with therapeutic failure). Over the course of the Milteforan™ therapy, the IC_50_ values of the isolates reached an average of 0.95 µM, approximately two times more resistant than the reference strain (considered sensitive) and much higher than the limit established for therapeutic success. This factor may further aggravate the limitations of Milteforan™-based monotherapy in dogs, which is already impaired in Brazil, among other factors, by the natural resistance of circulating parasites to this drug (41).

In addition, studies have shown that resistance to Milteforan™ remains constant even after passage through sandflies (42) and successive *in vitro* passages (39). This fact, combined with the wide use of Milteforan™ therapy to treat CVL in endemic areas, intense zoonotic transmission, and coupled with the fact that the dog can infect the invertebrate host even weeks after the end of treatment, despite being clinically cured (17), aggravate the problem involving the emergence of resistant parasites, since the dog can become infected with parasites that have already come into contact with the drug and, consequently, already present high resistance to it.

The isolates analyzed showed acquisition of resistance not only to the drug which they had contact with during the treatment of the dog (Miltefosine), but also to amphotericin B. This phenomenon, called cross-resistance, is well established in species of the genus *Leishmania* and involves several drugs(18,20,21,23,43,44). It is often associated with similar detoxification mechanisms by the parasite in response to different drugs, such as increased resistance to nitric oxide and expression of genes linked to the thiol metabolism of parasites exposed to meglumine antimoniate, which in turn also leads to increased resistance to allopurinol, another anti-leishmania drug with which metabolic protection routes are apparently shared (19,43,44). Although the detoxification mechanism shared between the drugs still remains uncertain, Mondelaers et al. (2018) reported clear cross-resistance between miltefosine and amphotericin B, corroborating the findings of the present study. In our analyses, a strong linear correlation (R2 = 0.83) was observed between the number of treatment courses with Milteforan™ to which the dog was submitted and the increase in the IC_50_ values of *L. infantum* isolates against amphotericin B. Acquisition of resistance to amphotericin B was similar to that to miltefosine, with IC_50_ values already higher than the control at T0, reaching values about 1.8 times higher after two courses of treatment with Milteforan™ (T2).

The presence of a cross-resistance phenomenon where the treatment of dogs with Milteforan™ generates parasites resistant not only to miltefosine, but also to amphotericin B extrapolates from the problem of the emergence of resistant parasites and goes beyond the scope of animal health, reaching human health, since this drug is one of the most commonly used to treat VL in humans. Thus, parasites resistant to both drugs could be easily transmitted to other dogs and, eventually, to humans due to the intense anthropozoonotic transmission in endemic areas and to the fact that resistance is maintained even after passage through sandflies (42). Amphotericin B, in its liposomal formulation, is used to treat VL in pregnant women, children under the age of one, or in individuals aged >50 years, with comorbidities, and who are HIV positive (45,46). All of these groups are considered at risk for the disease, requiring less toxic and more effective treatment, and this effectiveness that can be harmed by the emergence of parasites resistant to the drug.

Results of the impact of acquisition of resistance on the parasite fitness parameters did not show significant changes. The growth curve of the isolates did not change with increasing the number of treatment courses; however, these parasites showed significantly lower growth compared with that of the reference strain. This fact can be explained by a greater adaptation of the reference strain to *in vitro* growth, since it has remained for decades in this system. This factor is also possibly reflected in the production of metacyclic promastigote forms by isolated parasites in shorter cultivation times, since they have an optimized life cycle aiming at the infection of vertebrate hosts.

Results of the infectivity rates of the isolates against THP-1 cells demonstrated a reduction in the number of amastigotes per cell as the number of treatment courses to which the parasites were exposed increased. The current literature differs with respect to the impact of acquisition of drug resistance on parasite fitness. While some studies demonstrate that acquisition of resistance is followed by increased rates of infectivity, proliferation, and metacyclogenesis(24,25), other studies point out that the parasite has some of these parameters decreased in exchange for resistance, in a type of metabolic exchange currency (26,42,47,48), as it seems to occur in the present study in relation to the infectivity rate of the isolates.

Another parameter analyzed was the metacyclogenesis rate, regarding the production of metacyclic promastigote forms by the isolates. Although no statistical difference was observed between the different times analyzed, the metacyclogenesis rates show a clear upward trend as the number of treatment courses with Milteforan™ increases. This finding disagrees with what has been previously reported: acquisition of resistance leads to decreased parasite fitness. However, some authors consider the metacyclogenesis rate as one of the most important parameters to determine the virulence of a strain (27), since a larger number of infectious forms of *Leishmania* sp. represent greater spread of the disease in the host organism and even higher transmission rates between the vertebrate host and the invertebrate vector (49), which may further aggravate the finding in the present study involving the acquisition of resistance to two drugs used in animal and human therapy against VL.

Finally, it is worth mentioning that, although this study presents evaluations in isolated strains of only one dog, the nature of the research is unprecedented, since the same dog has been accompanied throughout its therapeutic process, enabling access to the same parasite at different stages of treatment and degrees of contact with the drug *in vivo*. Thus, it was possible to remove some biases from the study, such as genetic differences between different strains of the parasite, making it more robust, reliable, and contributing to the literature regarding the impact of treating dogs with Milteforan™ on the generation of resistant parasites.

## Conclusion

In conclusion, treating canine visceral leishmaniasis (CVL) with Milteforan™ induces *Leishmania infantum* resistance to miltefosine and amphotericin B in both forms of the parasite’s life cycle. Reduction in the infectivity rate with increasing the number of treatment courses in the dog, as well as a growth trend in the metacyclogenesis rates were observed. These factors can have a direct impact on the effectiveness of treating visceral leishmaniasis (VL) in animals and humans and, consequently, on public health.

## Acknowledgments

This study was funded by Conselho Nacional de Pesquisa e Desenvolvimento (CNPq) by the grant for productivity in research (309862/2015-9) and Professional Education Expansion Program (Proep) by the grant n° 442055/2019-6. This study also was supported by Instituto Carlos Chagas and Fundação Oswaldo Cruz (Fiocruz). The funders had no role in the decision to publish, or preparation of the manuscript. We declare no conflict of interest.

